# Neural dynamics of perceptual inference and its reversal during imagery

**DOI:** 10.1101/781294

**Authors:** Nadine Dijkstra, Luca Ambrogioni, Marcel A.J. van Gerven

## Abstract

After the presentation of a visual stimulus, cortical visual processing cascades from low-level sensory features in primary visual areas to increasingly abstract representations in higher-level areas. It is often hypothesized that the reverse process underpins the human ability to generate mental images. Under this hypothesis, visual information feeds back from high-level areas as abstract representations are used to construct the sensory representation in primary visual cortices. Such reversals of information flow are also hypothesized to play a central role in later stages of perception. According to predictive processing theories, ambiguous sensory information is resolved using abstract representations coming from high-level areas through oscillatory rebounds between different levels of the visual hierarchy. However, despite the elegance of these theoretical models, to this day there is no direct experimental evidence of the reversion of visual information flow during mental imagery and perception. In the first part of this paper, we provide direct evidence in humans for a reverse order of activation of the visual hierarchy during imagery. Specifically, we show that classification machine learning models trained on brain data at different time points during the early feedforward phase of perception are reactivated in reverse order during mental imagery. In the second part of the paper, we report an 11Hz oscillatory pattern of feedforward and reversed visual processing phases during perception. Together, these results are in line with the idea that during perception, the high-level cause of sensory input is inferred through recurrent hypothesis updating, whereas during imagery, this learned forward mapping is reversed to generate sensory signals given abstract representations.

---

When light hits the retina, a complex cascade of neural processing is triggered. Light waves are transformed into electrical signals that travel via the lateral geniculate nucleus of the thalamus to the visual cortex (1, 2). First, low-level visual features such as orientation and spatial frequency are processed in primary, posterior visual areas (3) after which activation spreads forward towards secondary, more anterior visual areas where high-level features such as shape and eventually semantic category are processed (4–6). This initial feedforward flow through the visual hierarchy is completed within 150 ms (7, 8) after which feedback processes are assumed to further sharpen representations over time until a stable percept is achieved (9, 10).

Activation in visual areas can also be triggered internally, in the absence of external sensory signals. During mental imagery, information from memory is used to generate rich visual representations. Neural representations activated during imagery are highly similar to those activated during perception (11). Imagining an object activates similar object representations in high-level visual cortex (12–15) and generating a mental image with simple visual features such as oriented gratings or letters is associated with perception-like activation of low-level visual areas (16–18).

In contrast, the temporal dynamics underlying the activation within the visual system during perception and mental imagery are presumably very different. The neural dynamics of the early stages of perception have been extensively characterized with intracranial electrophysiological recordings in primates (3, 5, 8). However, the neural dynamics of imagery, i.e. how activation travels through the brain during internally generated visual experience, remain unclear. Researchers from various fields have proposed that the direction of information flow during internally generated visual experience is reversed compared to perception (19–22). In line with this idea, a recent study showed that during memory recall, high-level, semantic representations were active before low-level, perceptual representations (23). However, the localization of activation in this study was ambiguous such that it is possible that all processing happened within high-level visual cortex but that only the dimension to which the neurons were sensitive changed from abstract features to perceptual features over time (for an example of dynamic neural tuning, see (24)). Moreover, memory recall was not directly compared with memory encoding. Therefore, it remains unclear whether the same perceptual cascade of neural activation is reactivated in reverse order during internally generated visual experience.

According to predictive processing (PP) theories, reversals of information flow also play an important role during perception. PP states that the brain deals with the inherent ambiguity of incoming sensory signals by incorporating prior knowledge about the world (25). This knowledge is used to generate top-down sensory predictions which are compared to the bottom-up sensory input. Perceptual inference is then accomplished by iteratively updating the model of the world until the difference between prediction and input is minimized (26, 27). Therefore, the neural dynamics of stimulus information during perception should be characterized by an interplay between feedforward and feedback sweeps. Simulations based on PP models predict that these recurrent dynamics are dominated by slow-wave oscillations (28, 29).

In this study, we used magnetoencephalography (MEG) and machine learning to characterize the spatio-temporal dynamics of information flow during mental imagery and perception. We first characterized neural activity during the initial perceptual feedforward sweep using multivariate classifiers at different time points which served as proxies for representations in different visual areas. That is, decoding at early perception time points was taken to reflect stimulus representations in low-level, posterior visual areas while decoding at later time points was taken to reflect high-level, anterior visual representations. Then, we estimated when these feedforward perception models were reactivated during imagery and later stages of perception. Our results reveal that, while the neural stimulus representations activated during perception and imagery are similar, the temporal dynamics underlying this activation is very different, revealing a fundamental asymmetry in neural information processing.

## Results

Twenty-five participants executed a retro-cue task while MEG was measured. During the task, two consecutive stimuli were presented, a face and a house or a house and a face, followed by a cue indicating whether participants had to imagine the first or the second stimulus. They then imagined the cued stimulus as vividly as possible and indicated their experienced imagery vividness. To ensure that participants were generating detailed mental images, we included catch trials during which participants had to indicate which of four highly similar exemplars they had just imagined. An accuracy of 89.9 % (*SD* = 5.4%) indicated that participants did indeed imagine the stimuli with a high degree of visual detail.

### Inferring information flow using perceptual feedforward classifier models

Classifier models representing neural representations at different time points during perception were obtained using linear discriminant analysis (LDA). An LDA classifier was trained to decode the stimulus class (‘face’ vs ‘house’) from sensor level activity at each time point, giving different perception models for different time points (Fig. 1A). We decided to only focus on the time period between 70 ms and 130 ms after stimulus onset, which, with a sampling rate of 300 Hz, contained 19 perception models. Perceptual content could be decoded significantly better than chance level at 70 ms after stimulus onset (30). This is in line with intracranial studies which showed that visual information is detectable in early visual cortex from 50 ms onwards (5). Furthermore, both intracranial as well as scalp electrophysiology has shown that around 130 ms, high-level object representations first get activated (4, 6, 7, 31). Therefore, this early time window is representative of the feedforward sweep during perception. In line with this, source reconstruction of the sensor-level activation patterns at different time points shows that stimulus information spreads from low-level visual areas towards higher-level visual areas during this period (Fig. 1A). Furthermore, cross-correlation between the information flow in early visual cortex (EVC) and inferior temporal cortex (IT; Fig. 1B) confirms that stimulus information is available in low-level EVC 26.3 ms (*CI* = 13.1 to 32.9 ms) before it reaches high-level IT.

**Fig. 1.**
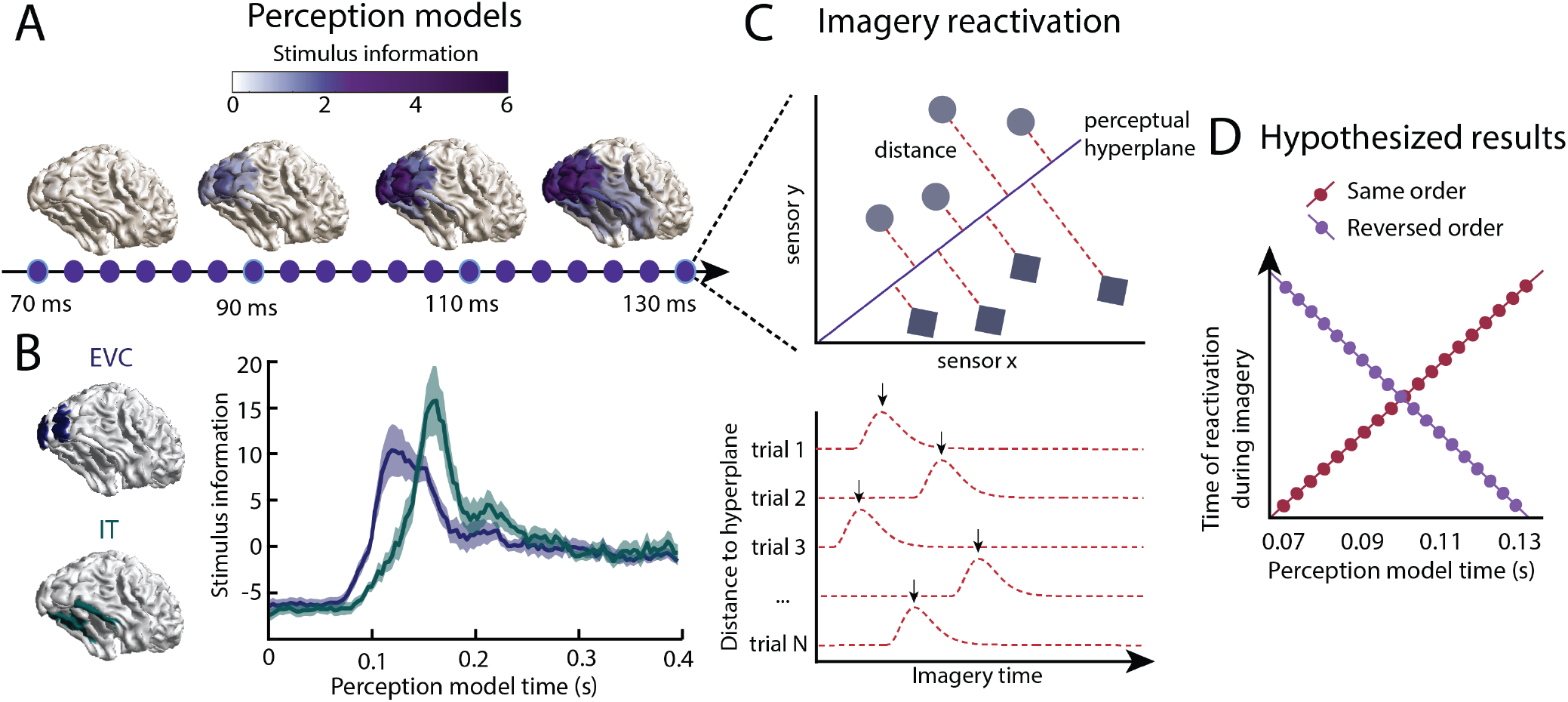
Inferring information flow using perceptual feedforward classifier models. (A-B) Perception models. at each point in time between 70 and 130 ms after stimulus onset, a perception model (classifier) was estimated using Linear Discriminant Analysis (LDA) on the activation patterns over sensors. (A) The source-reconstructed difference in activation between faces and houses (i.e. decoding weights or stimulus information) is shown for different time points during perception. (B) The normalized stimulus information over time is shown for low-level early visual cortex (EVC: blue) and high-level inferior temporal cortex (IT: green). These data confirm a feedforward flow during the initial stages of perception. (B) Imagery reactivation. For each trial and time point during imagery, the distance to the perceptual hyperplane of each perception model is calculated. The timing of the reactivation of each during imagery is determined by finding the peak distance for each trial. (C) Hypothesized results. This procedure results in a measure of the imagery reactivation time for each trial, for each perception model time point. If perception models are reactivated in the same order during imagery, there would be a positive relation between reactivation imagery time and perception model time. If instead, perception models are reactivated in reverse order, there would be a negative relation.

To identify when these representations were reactivated during imagery, we tested the perception models on imagery to obtain the distances to the classifier hyperplanes per trial (Fig. 1C). The distance to the hyperplane indicates the amount of classifier evidence present in the data. Distance measures have previously been used as a measure of model activation (23, 32) and have been linked to reaction time measurements (33). For each perception model and for each imagery trial, we identified the time of the absolute peak distance (23). This resulted in a trial-by-trial estimate of the reactivation timing for the different perception models. If processing happens in a similar order during imagery as during perception, we would expect that during imagery, early perception models are reactivated earlier in time than late perception models (Fig. 1D). This would result in a positive relation between perception model time and imagery reactivation time. If instead, processing happens in reverse order, with late, high-level models being active before earlier models, we would see a negative relation between perception model time and imagery model time.

### Perceptual feedforward sweep is reversed during imagery

The reactivation time during imagery for the different perception models is shown in Figure 2A. To test whether there was a significant ordering in the reactivation we ran a linear mixed-effects model (LMM) with the reactivation time during imagery as dependent variable, the perception model time as fixed predictor, and subject and trial as random variables. Five models with different combinations of random effects were estimated and the model with the highest Schwarz Bayesian Information Criterion (BIC) was used to ensure best-fit with minimum number of predictors (Supplementary Table S1). The winning model contained a random effect for the intercept of each subject and trial. This means that this model allowed the reactivation of the perception sequence to start at different time points per trial and per subject, which is in line with the idea that there is a large variation in timing of imagery between trials and subjects (30).

**Fig. 2.**
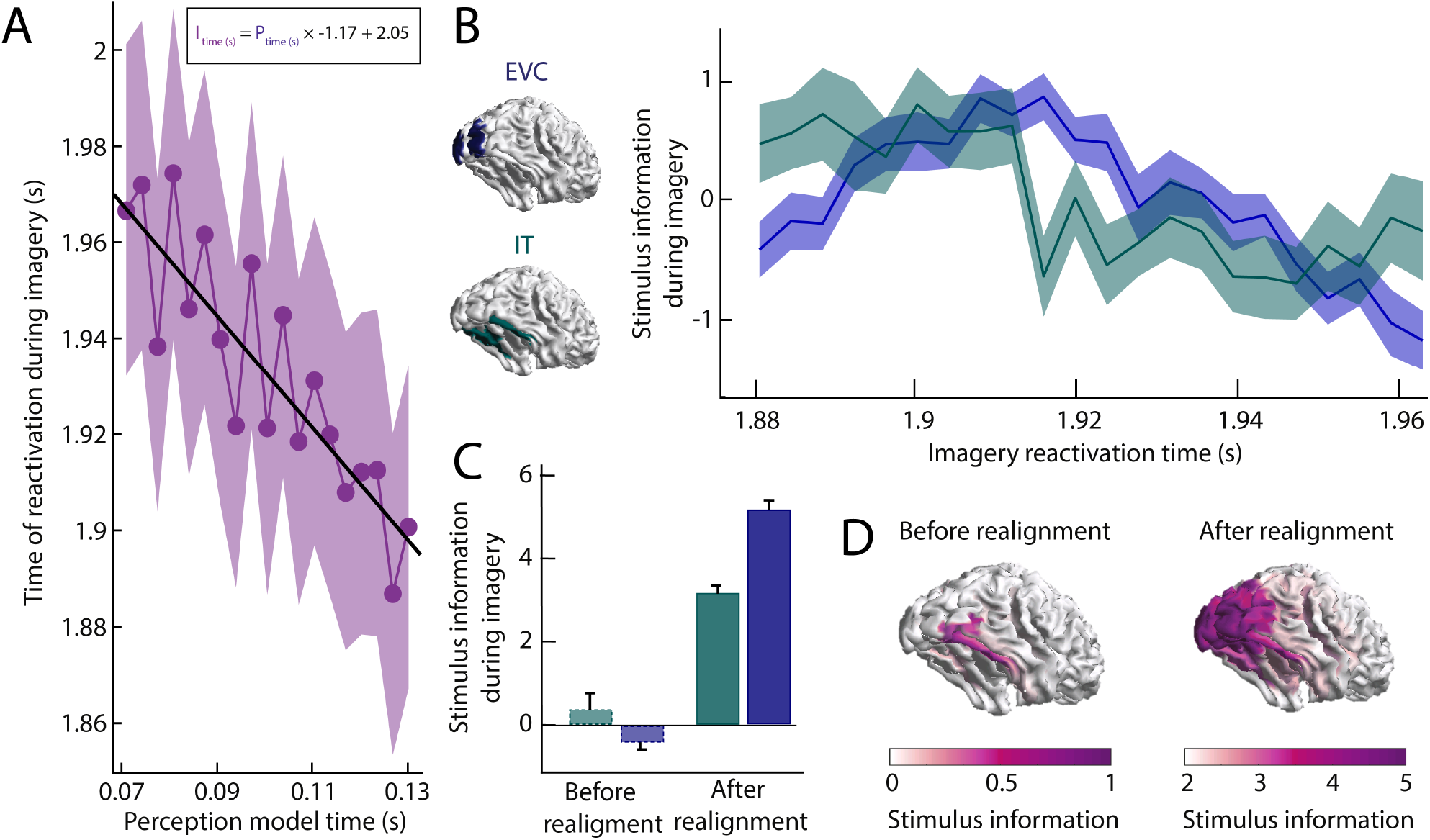
Imagery reactivation results. (A) Reactivation time during imagery for every perception model averaged over trials. The shaded area represents the 95% confidence interval. The linear equation shows how imagery reactivation time (I) can be calculated using perception model time (P) in seconds. (B-D) Stimulus information during imagery was estimated by realigning the trials based on the reactivation time points and using the linear equation to estimate the imagery time axis. (B) Normalized stimulus information over time for low-level early visual cortex (EVC) and high-level inferior temporal cortex (IT). (C) Stimulus information in EVC and IT averaged over time before and after realignment. (D) Source reconstruction of stimulus information averaged over time before realignment around peak time-locked imagery decoding (left) and after realignment (right). Stimulus information below 0 indicates that the amount of information did not exceed the permutation distribution.

The model showed a significant main effect of perception time (*t*(98160) = −5.403, *p* = 6.58e-8) with a negative slope (*β*0 = 2.047, SD = 0.282; *β*1 = −1.171, SD = 0.217) indicating that models of later perception time points were associated with earlier imagery reactivation times. The fact that the absolute slope is so close to one suggests that reactivation during imagery of the perception model sequence happens at a similar speed as the original activation during perception. Next, we reconstructed the imagery activation by realigning the trials based on the identified peak time points for each perception model time point. The imagery time line was inferred using the linear equation obtained from the LMM. The temporal dynamics of high-level IT and low-level EVC (Fig. 2B) confirm the conclusion that during imagery, information flows from high-level to low-level visual areas. Crosscorrelation between these realigned signals shows that information in IT precedes information in EVC by 11.2 ms (*CI* = 0 to 29.9 ms). Furthermore, before realignment, time-locked decoding during imagery only revealed a small amount of information in high-level visual cortex (Fig 2D). In contrast, after realignment, stimulus information was clearly present throughout the entire visual hierarchy. The net increase for low and high-level regions is shown in Figure 2C: before realignment, there was no measurable information in low-level EVC during imagery, but after realignment there clearly was. This emphasizes how time-locked analyses obscure neural processing during complex cognitive processes such as mental imagery.

To ensure that these results were not due to confounds in the structure of the data irrelevant to reactivation of stimulus representations, we performed the same analysis after permuting the labels of the models. Specifically, we trained the perception models using random class assignments and then again calculated the reactivations during imagery. The results are shown in Supplementary Figure S1. The sequential reactivation disappeared when using shuffled classifiers as perception time did not significantly predict imagery reactivation time anymore (*t*(98160) = −0.762, *p* = 0.446). Furthermore, for the main analysis, we removed high frequency noise from the imagery distance traces. To check whether this filter somehow altered the results, we ran the same analysis without the low-pass filter, which gave similar results (Supplementary Figure S2).

### Reactivation during imagery reveals recurrent processing during perception

In the previous analysis we focused on the first 150 ms after stimulus presentation because this period reflects the initial perceptual feedforward sweep. A negative relationship with imagery reactivation time therefore indicated feedback processing during imagery (cf. Fig. 2). However, feedback processes are assumed to play a fundamental role in later stages of perception (34, 35). For these later stages we would therefore expect a positive relation with imagery reactivation, indicating that information flows in the same direction. To investigate this, we next calculated the imagery reactivation time for all time points during perception (Fig. 3A). The results between 70 ms and 130 ms are equivalent to Figure 2A.

**Fig. 3.**
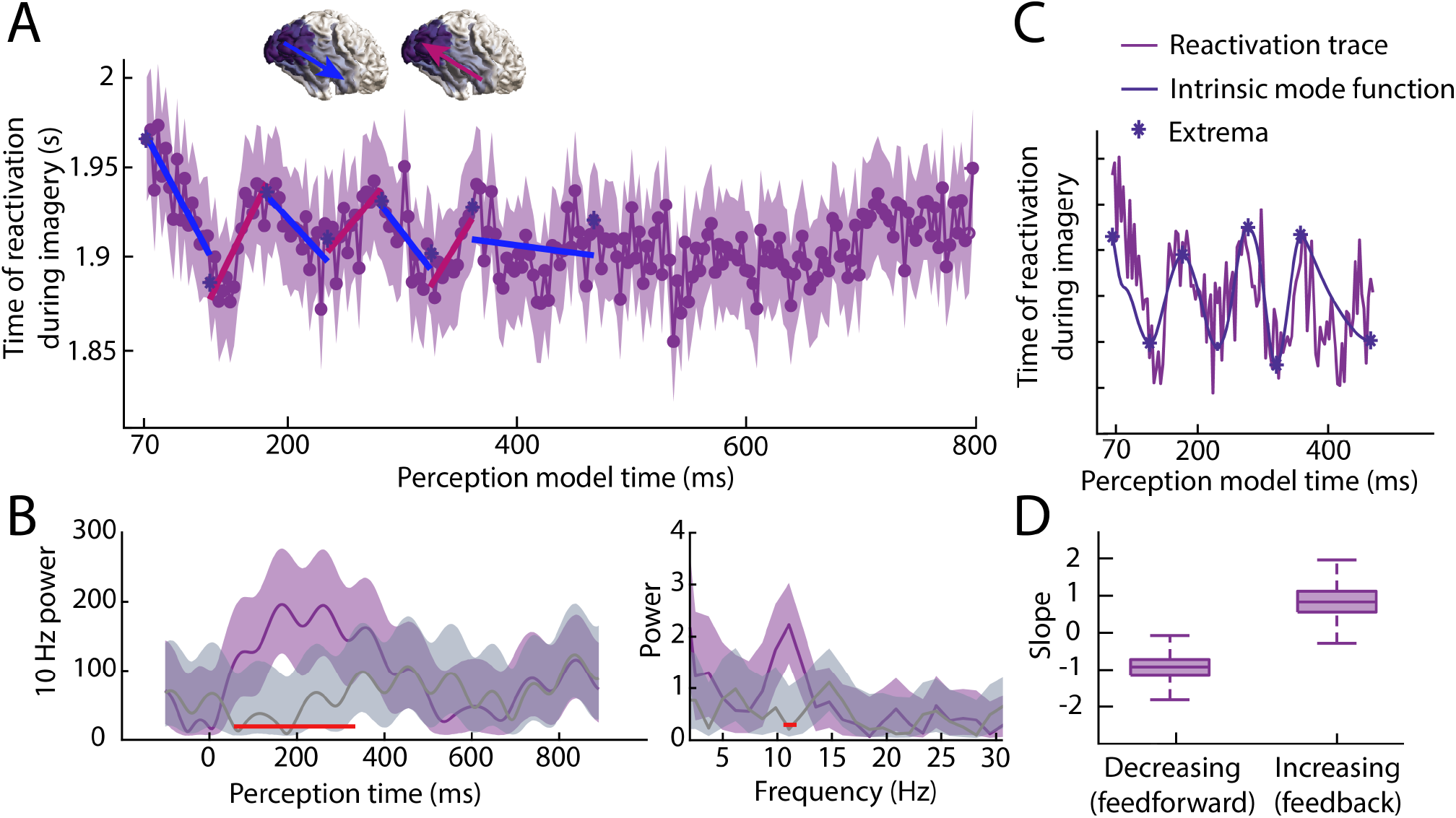
Reactivation timing during imagery for classifiers trained at all perception time points. (A) Imagery reactivation time for perception models trained on all time points. On the x-axis the training time point during perception is shown and on the y-axis the reactivation time during imagery is shown. The dots represent the mean over trials for individual time points and the shaded area represents the 95% confidence interval. (B) Left: time-frequency decomposition using a Morlet wavelet at 10 Hz. Right: power at different frequencies using a Fast Fourier Transformation. The purple line represents the true data and the grey line represents the results from the shuffled classifier. Shaded areas represent 95% confidence intervals over trials. Red lines indicate time points for which the true and shuffled curve differed significantly (FDR corrected). (C) Intrinsic mode function and its extrema derived from the reactivation traces using empirical model decomposition. (D) Average decreasing and increasing slopes, boxplots reflect uncertainty over bootstrapping samples.

Interestingly, during the first 400 ms of perception, there seems to be an oscillatory pattern in the relationship between perception time and reactivation time during imagery, where positive and negative slopes alternate. This pattern repeats four times in 400 ms, roughly reflecting an alpha oscillation. To investigate this further, we quantified 10 Hz power over time using a Morlet decomposition (Fig. 3B left, purple curve) and compared the results with the permuted classifier (Fig. 3B left, cf. Fig. S1). There was a significant increase in 10 Hz power between 80 and 315 ms after stimulus onset (all FDR corrected *p*-values below 0.003). Furthermore, a Fast Fourier Transform over the first 400 ms revealed that the difference in power was limited to the 11.2 Hz frequency (Fig. 3B right, *p* = 0.001).

These results indicate that, during perception, stimulus information travels up and down the visual hierarchy aligned to the alpha frequency. We next aimed to investigate the speed of feedforward and feedback sweeps by looking at the absolute slopes of the increasing and decreasing phases. To this end, we used empirical mode decomposition (EMD) over the first 400 ms to seperate the signal in intrinsic mode functions (IMFs). We selected identified increasing and decreasing phases by selecting the period between two subsequent extrema (Fig. 3C) and calculated the slope for each phase (Fig. 3D, single estimated slopes are shown in Fig. 3A). The average decreasing (feedforward) slope was −0.69 (*CI* = −2.05 to −0.35) and the average increasing (feedback) slope was 1.02 (*CI* = −0.31 to 2.14; Fig. 3D). There was no significant difference in the absolute slope value between increasing and decreasing phases (Mdiff = 0.33, *CI:* −1.80 to 1.74), revealing no evidence for a difference in processing speed for feedforward and feedback sweeps.

It is possible that the oscillation in the reactivation trace is due to evoked alpha oscillations in the raw signal which modulates signal-to-noise ratio via its amplitude. To control for this, we calculated the spectral coherence at 11.2 Hz between the reactivation signal and the raw data signal at each sensor over subjects. We then compared the coherence between the raw data and the true reactivation with the coherence between the raw data and the shuffled reactivation using a permutation test with 10000 permutations. None of the sensors showed a significant difference in coherence between the true and permuted classifier (FDR corrected), even though the reactivation oscillation was only present for the true classifier (cf. Fig 3B). This indicates that the oscillatory pattern in the reactivation trace does not just reflect evoked alpha in the raw signal.

### Stimulus representations are iteratively updated during perception

If there is indeed a recurrent information flow up and down the visual hierarchy during perception, we should also be able to demonstrate this within perception. The previous results suggest specific time windows of feedforward and feedback phases during perception. To test whether these different phases indeed reflected reversals of information flow, we applied the reactivation analysis previously used for imagery (cf. Fig. 1) to the different perception phases identified in the previous analysis (cf. Fig. 3). We predicted that perception classifiers trained during a decreasing (feedforward) phase would be reactivated in reverse order during an increasing (feedback) phase and vice versa, showing a negative relationship between training time and reactivation time. In contrast, classifiers trained and tested on the same type of phase (both decreasing or both increasing), should show a positive relationship (see Fig. 4B, for hypothesized results).

**Fig. 4.**
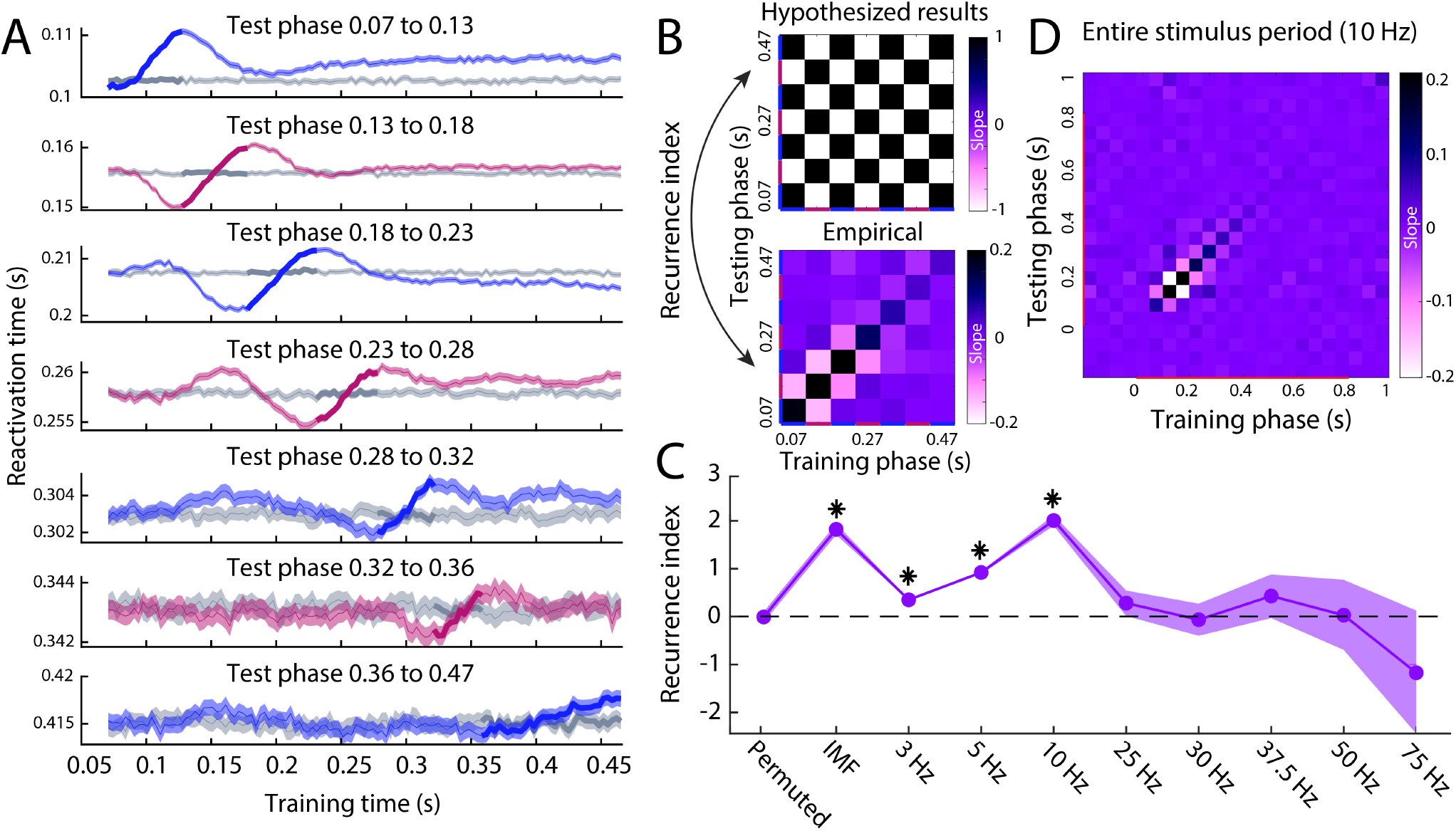
Reactivation timing for different perception phases. (A) The reactivation traces for each testing phase. Blue traces reflect feedforward phases, pink traces reflect feedback phases (cf. Fig. 3A) and grey traces reflect reactivation traces for permuted classifiers. Shaded area represents the 95% confidence interval over trials. (B) Hypothesized (top) and empirical (bottom) slopes between the training and testing phases. The hypothesized matrix assumes recurrent processing such that subsequent phases show a reversal in the direction of information flow. Recurrence index reflects the amount of recurrent processing in the data which is quantified as the dot product between the vectorized hypothesis matrix and empirical matrix. (C) Recurrence index for the permuted classifier, phase specification based on the IMF of the imagery reactivation trace (cf. Fig. 3C) and phase specification on evoked oscillations at various frequencies. (D) Slope matrix for phase specification defined at 10 Hz over the entire stimulus period.

The reactivation traces for the different test phases are shown in Figure 4A. Blue traces represent reactivations during feedforward phases and pink traces represent reactivations during feedback phases. Grey traces show the results for a permuted classifier. In line with the previous findings, for most phases, there is a clear oscillatory pattern between training time and reactivation time within perception. For each phase, the training time corresponding to that testing phase is highlighted in bold. This time period should always show a positive slope, indicating that classifiers trained and tested on time points belonging to the same phase are reactivated in the same order.

The slopes between the training time and the reactivation time of the different phases are shown below the hypothesized results in Figure 4B. As expected, the diagonal, reflecting training and testing on the same phase, was always positive. In contrast, training and testing on different phases tended to be associated with negative slopes. Note that for all decoding analyses, cross-validation was used, which means that these results cannot be due to overfitting but reflect true representational overlap between phases. To quantify the effect, we calculated a *Recurrence Index* (RI) which was defined as the dot-product between the vectorized hypothesis-matrix and the empirical-matrix. The RI is positive if the data shows the hypothesized oscillatory pattern of slope reversals, zero if there is no clear oscillatory pattern and negative if the data shows the opposite pattern. The RI was significantly larger than zero for the true data (RI = 1.83, *CI* = 1.71 to 1.94, *p* < 0.0001) but not for the permuted data (RI = −0.008, *CI* = −0.12 to 0.10, *p* = 0.548). This confirms that during perception, stimulus information flows up and down the visual hierarchy in feedback and feedforward phases.

The phases that we used here were identified based on the IMF of the oscillation in the imagery reactivation trace (cf. Fig. 3), leading to phases of different lengths. Next, we investigated whether we could observe the same pattern if we specified the phases based on fixed evoked oscillations at different frequencies. For example, a 10 Hz oscillation resulted in 4 feedforward and 4 feedback phases of 50 ms each within our 400 millisecond time window. The RI for the different frequencies is plotted in Figure 4C. Whereas the RI was significantly above zero for several low frequencies, the oscillatory pattern was clearest for the IMF based phases and for the 10 Hz oscillation, confirming that the perceptual recurrence is most strongly aligned to the alpha frequency. Furthermore, to investigate whether this pattern of recurrence was indeed specific to the 400 millisecond time window identified previously, we also applied the 10 Hz recurrence analysis to the entire stimulus period (Fig. 4D). The results show that the recurrence pattern is indeed restricted the first 400 ms after stimulus onset.

An interesting observation is that the recurrence pattern seems to be restricted to around the testing phase, such that only classifiers trained on phases close to the testing phase show a clear positive or negative relation with reactivation time. This is what causes the ‘traveling wave’ pattern between the rows in Figure 4A. An intriguing explanation for this observation is that stimulus representations change over subsequent cycles, such that representations only show a reactivation relation with neighboring phases, but that the representations during later phases are too dissimilar to result in reliable reactivations. We tested whether recurrence was indeed specific to phases around the testing phase by comparing the normalized RI of slopes next to the diagonal in the slope matrix with the other slopes. Neighboring phases indeed show a significantly higher RI (*M* = 0.073, *CI* = 0.069 to 0.076) than other phases (*M* = 0.002, *CI* = −0.0004 to 0.005, *p* < 0.0001), confirming that recurrence was restricted to a small number of cycles.

## Discussion

In this study we investigated how stimulus representations flow through the visual system during mental imagery and perception. Our results reveal an asymmetry in information processing between perception and imagery. First, we showed that early perception processes are reactivated in reverse order during imagery, demonstrating that during imagery, activation flows from high-level visual areas down to low-level areas over time. Second, for later stages of perception, we found an oscillatory pattern of alternating positive and negative relations with imagery reactivation, indicating recurrent stimulus processing up and down the visual hierarchy aligned to an 11Hz oscillation. Finally, by focusing on the identified feedforward and feedback phases, we showed that recurrence during perception was restricted to neighbouring phases, suggesting that the format of neural stimulus representations changed with subsequent cycles of recurrent processing. Together, these findings indicate that during imagery, stimulus representations are activated via feedback processing whereas during perception, stimulus representations are iteratively updated through cycles of recurrent processing.

Our results are neatly in line with predictive processing (PP) theories. According to PP, recurrent processing during perception reflects dynamic hypothesis testing (26–28, 36). Specifically, perceptual inference is assumed to be accomplished via an interplay between top-down prediction signals encoding perceptual hypotheses and bottom-up prediction errors encoding the sensory signal unexplained by these hypotheses. Inferring the cause of sensory input is done by iteratively updating the perceptual hypothesis until the prediction error is minimized, in line with the dynamically changing representations observed here. Importantly, recurrent processing is assumed to happen hierarchically such that each level is activated by both bottom-up evidence as well as top-down predictions (26). This is in line with our observation that feedforward and feedback sweeps proceeded at the same speed. Also in line with the current findings, PP predicts that these recurrent dynamics are dominated by slow-wave oscillations (28). To our knowledge, the current study is the first to show these perceptual updating cycles empirically in humans. Furthermore, our results suggest that in the current task context, the perceptual inference process was completed in approximately four updating cycles. An exciting avenue for future research is to investigate whether the number of cycles needed can be modulated by task variables such as attention and stimulus noise.

Whereas perception was characterized by dynamically changing representations updated through recurrent cycles, we only found evidence for a single feedback flow during imagery. Specifically, all perceptual feedforward sweeps showed a negative relationship within the same imagery time window and all perceptual feedback sweeps showed a positive relationship within that same imagery time window, suggesting that the imagery feedback flow contained the complete stimulus representation that was inferred during perception. This result fits with the idea that imagery uses the same predictive processes that underlie perceptual inference to run off-line simulations of sensory representations under different hypotheses (11, 37–40). In contrast to perception, during imagery, there is no bottom-up sensory input prompting hypothesis updating. Instead, the perceptual cause is given and feedback connections are used to generate the corresponding low-level sensory representation based on the mapping that was learned during perception. Recurrent processing within the visual system might become important when imagining a more dynamic stimulus in which sensory representations change over time. However, it is also possible that recurrent dynamics were actually present during imagery in this task but that we were unable to reveal them due to signal-to-noise issues. Future research should focus on developing more sensitive techniques to further characterize information flow during imagery. Another interesting question for future research is whether reactivation during mental imagery has to always fully progress down the visual hierarchy. In this study, participants were instructed to generate highly detailed mental images and catch trials were used to ensure that they indeed focused on low-level visual details. It might be the case that if less detail is needed for the task, earlier perception processes are not reactivated and mental simulations stop at a higher level (22, 41).

In conclusion, by generalizing multivariate decoding models between brain activity at different time points, we show that, while similar neural representations are activated during imagery and perception, the neural dynamics underlying this activation are different. Whereas perception is characterized by recurrent processing, imagery is dominated by top-down feedback processing. These results are in line with the idea that during perception, high-level causes of sensory input are inferred whereas during imagery, this inferred mapping is reversed to generate sensory representations given these causes. This highlights a fundamental asymmetry in information processing between perception and imagery and sets the stage for exciting new avenues for future research.

## Materials and methods

### Participants

Thirty human volunteers with normal or corrected-to-normal vision gave written informed consent and participated in the study. Five participants were excluded: two because of movement in the scanner (movement exceeded 15 mm), two due to incorrect execution of the task (less than 50% correct on the catch trials, as described below) and one due to technical problems. Twenty-five participants (mean age 28.6, SD = 7.62) remained for the final analysis. The study was approved by the local ethics committee and conducted according to the corresponding ethical guidelines (CMO Arnhem-Nijmegen). An initial analysis of these data has been published previously (30).

### Experimental design

We adapted a retro-cue paradigm in which the cue was orthogonalized with respect to the stimulus identity (42). Participants were shown two images after each other followed by a retro-cue indicating which of the images had to be imagined. After the cue, a frame was shown in which the participants had to imagine the cued stimulus as vividly as possible. Next, they had to indicate their experienced imagery vividness by moving a bar on a continuous scale. To ensure that participants were imagining the stimuli with great visual detail, both categories contained eight exemplars, and on 7% of the trials the participants had to indicate which of four exemplars they imagined. The exemplars were chosen to be highly similar in terms of low-level features to minimize within-class variability and increase between-class classification performance. We instructed participants to focus on vividness and not on correctness of the stimulus, to motivate them to generate a mental image including all visual features of the stimulus. The stimuli encompassed 2.7 x 2.7 visual degrees. A fixation bull’s-eye with a diameter of 0.1 visual degree remained on screen throughout the trial, except during the vividness rating.

### MEG recording and preprocessing

Data were recorded at 1200 Hz using a 275-channel MEG system with axial gradiometers (VSM/CTF Systems, Coquitlam, BC, Canada). For technical reasons, data from five sensors (MRF66, MLC11, MLC32, MLF62, MLO33) were not recorded. Subjects were seated upright in a magnetically shielded room. Head position was measured using three coils: one in each ear and one on the nasion. Throughout the experiment head motion was monitored using a real-time head localizer (43). If necessary, the experimenter instructed the participant back to the initial head position during the breaks. This way, head movement was kept below 8 mm in most participants. Furthermore, both horizontal and vertical electrooculograms (EOGs), as well as an electrocardiogram (ECG) were recorded for subsequent offline removal of eye- and heart-related artefacts. Eye position and pupil size were also measured for control analyses using an Eye Link 1000 Eye tracker (SR Research). Data were analysed with MATLAB version R2018a and FieldTrip (44) (RRID: SCR_004849). Per trial, three events were defined. The first event was defined as 200 ms prior to onset of the first image until 200 ms after the offset of the first image. The second event was defined similarly for the second image. Further analyses focused only on the first event, because the neural response to the second image is contaminated by the neural response to the first image. Finally, the third event was defined as 200 ms prior to the onset of the retro-cue until 500 ms after the offset of the imagery frame. As a baseline correction, for each event, the activity during 300 ms from the onset of the initial fixation of that trial was averaged per channel and subtracted from the corresponding signals. The data were downsampled to 300 Hz to reduce memory and CPU load. Line noise at 50 Hz was removed from the data using a DFT notch filter. To identify artefacts, the variance of each trial was calculated. Trials with high variance were visually inspected and removed if they contained excessive artefacts. After artefact rejection, on average 108 perception face trials, 107 perception house trials, 105 imagery face trials and 106 imagery house trials remained for analysis. To remove eye movement and heart rate artefacts, independent components of the MEG data were calculated and correlated with the EOG and ECG signals. Components with high correlations were manually inspected before removal. The eye tracker data was cleaned separately by inspecting trials with high variance and removing them if they contained blinks or other excessive artefacts.

### Temporal decoding analysis

To estimate when neural representations at different perception time points got reactivated during imagery, we trained classifiers on several time points during perception and applied them to imagery. Specifically, to decode the stimulus category per perception time point, we used a linear discriminant analysis (LDA) classifier with the activity from the 270 MEG sensors as features (see (45) for more details). To prevent a potential bias in the classifier, the number of trials per class was balanced per fold by randomly removing trials from the class with the most trials until the trial numbers were equal between the classes. In our design, the perception and imagery epochs happened in the same general ‘trial’. If the imagery epochs on which the classifier was tested came from the same trials as the perception epochs on which it was trained, auto-correlations in the signal could inflate decoding accuracy. To circumvent this, a five-fold cross-validation procedure was implemented where for each fold the classifier was trained on 80% of the trials and tested on the other 20%. Before decoding the imagery signal, the mean perception signal over trials per sensor was subtracted to increase sensitivity to changes in multivariate patterns instead of amplitude.

This resulted in a decision value for each perception model, for each imagery trial and time point. The sign of this value indicates in which category the trial has been classified and the value indicates the distance to the hyperplane. In order to obtain the confidence of the classifier in the correct class, the sign of the distance values in one category was inverted. This means that increasing positive distance values now always reflected increasingly confident classification. To obtain the specific moment within each imagery trial that a given perception model became active, we identified the time point with the highest distance (23) (cf. Fig. 1).

After discovering the oscillatory pattern between later perception time points and imagery reactivation, we also ran this reactivation analysis within perception by training on all perception time points and testing on specific perception time windows reflecting the identified feedforward and feedback phases (cf. Fig. 3). This identified, per phase, reversals in information flow.

For the imagery generalization, the time window used to obtain the peak distance extended from the cue onset until the vividness instruction onset, covering the entire 4 seconds during which participants were instructed to imagine the stimulus. For this data, we removed high frequency noise using a low-pass filter of 30 Hz (for the results using the raw data, see Supplementary Figure S2). For the within-perception tests we did not use a low-pass filter because we used smaller testing time windows and were also interested in possible high-frequency effects.

### Realignment

To further investigate the effects of trial by trial differences in imagery timing on stimulus representations, we realigned the imagery activation based on the identified distance peaks. This was done by selecting the peak imagery time point for every trial for every perception model time point to create a realigned imagery data set. The time axis for this new data set was inferred using the linear relationship between perception model time and imagery reactivation time established in the main imagery reactivation analysis. Source activation was then calculated using the same procedure as was used for the un-realigned perception and imagery data (more details below under *Source Localization*).

### Frequency analysis

For the time frequency analysis, we used a Morlet Wavelet at 10 Hz defined as:

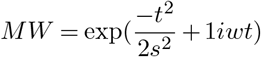

where *t* is a time vector from −0.5 to 0.5 in steps of 1/fs, *w* = 2π10, 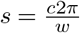 and *c* is the number of cycles, this case 1. We only used 200 samples in the centre of the wavelet and convolved this with the mean reactivation trace to obtain the time frequency representation. To calculate power at different frequencies we used the Fast Fourier Transform (FFT). To prevent edge effects, we first multiplied the mean signal with a Hanning taper from −0.2 to 0.6 seconds prior to performing the FFT. Of the resulting complex numbers, the absolute value was taken and the result was normalized by the length of the signal.

Within reactivation traces, positive slopes represented reactivation in a similar order whereas negative slopes represented reactivation in reverse order. We wanted to divide the signal into phases of positive and negative slopes because these represented feedforward and feedback phases. In order to do this, we used empirical mode decomposition (EMD) which separates the signal into intrinsic mode functions (IMF) based on local and global extrema; i.e. peaks and troughs (46–48). For a 10 Hz frequency we would expect eight extrema in 400 ms, reflecting four full cycles. Therefore, to identify the different phases, we selected the IMF with the number of extrema closest to eight. Decreasing and increasing phases were defined as periods between subsequent extrema (cf. Fig. 3C). Slopes between periods were calculated using linear regression and slopes for decreasing and increasing periods were averaged to reflect the speed of feedforward and feedback processing respectively. To determine the uncertainty of these slopes, for every bootstrapping sample the mean reactivation trace and the corresponding EMD separation was recalculated.

### Statistics

To test whether there was a significant linear relationship between perception training model time and imagery reactivation time, we used a generalised linear mixed model (GLMM) with the single trial classifier distance peaks as dependent variable and perception model time during the feedforward sweep as independent variable. We chose GLMMs because they make fewer assumptions than more commonly used GLMs and because we expected large differences in the onset of reactivation between trials and subjects and, in contrast to GLMs, GLMMs allow for random effects on trials and subjects.

To obtain 95% confidence intervals for reactivation times, time-frequency and frequency plots, we performed bootstrapping analyses with 10000 bootstrapping samples. For pairwise comparisons, we obtained p-values by bootstrapping the difference between the two conditions. Source traces represented the mean difference between the stimulus classes which cannot be computed per trial. Therefore, uncertainty in the mean of these values was represented as the standard error of the mean (SEM) over subjects.

### Source localization

To identify brain areas that represented information about the stimuli during perception and imagery, we performed source reconstruction. For LDA classification, the spatial pattern that underlies the classification (i.e. the decoding weights), reduces to the difference in magnetic fields between the two conditions (49). Therefore, the difference ERF between faces and houses reflects the contributing brain areas. For the sensor-level activation plots, we calculated the planar gradient for each participant prior to averaging over participants. For the source-level plots, we performed source reconstruction on the axial difference ERF. T1-weighted structural MRI images were acquired in separate sessions using a Siemens 3T MRI scanner. Vitamin E markers in both ears indicated the location of the head coils during the MEG measurements, allowing for realignment between the two. The location of the fiducial at the nasion was estimated based on anatomy. The volume conduction model was created based on a single shell model of the inner surface of the skull. The source model was based on a reconstruction of the cortical surface created for each participant using FreeSurfer’s anatomical volumetric processing pipeline (RRID: SCR_001847). MNE-suite (Version 2.7.0; RRID: SCR_005972) was subsequently used to infer the subject-specific source locations from the surface reconstruction. The resulting head model and source locations were co-registered to the MEG sensors.

The lead fields were rank reduced for each grid point by removing the sensitivity to the direction perpendicular to the surface of the volume conduction model. Source activity was obtained by estimating linearly constrained minimum variance (LCMV) spatial filters (50). The data covariance was calculated over the interval of 50 ms to 1 s after stimulus onset for perception and over the entire segment for imagery. The data covariance was subsequently regularized using shrinkage with a regularization parameter of 0.01 (as described in (51)). These filters were then applied to the sensor MEG data, resulting in an estimated two-dimensional dipole moment for each grid point over time.

To facilitate interpretation and visualization, we reduced the two-dimensional dipole moments to a scalar value by taking the norm of the vector. This value reflects the degree to which a given source location contributes to activity measured at the sensor level. However, the norm is always a positive value and will therefore, due to noise, suffer from a positivity bias. To counter this bias, we employed a permutation procedure in order to estimate this bias. Specifically, in each permutation, the sign of half of the trials were flipped before averaging and projecting to source space. This way, we cancelled out the systematic stimulus-related part of the signal, leaving only the noise. Reducing this value by taking the norm thus provides an estimate of the positivity bias. This procedure was repeated 1000 times, resulting in a distribution of the noise. We took the mean of this distribution as providing the most likely estimate of the noise and subtracted this from the true, squared source signal. Furthermore, this estimate provides a direct estimate of the artificial amplification factor due to the depth bias. Hence, we also divided the data by the noise estimate to obtain a quantity that allowed visualization across cortical topography. Values below zero therefore reflected no detectable signal compared to noise. For full details, see (51). To perform group averaging, for each subject, the surface-based source points were divided into 74 atlas regions as extracted by FreeSurfer on the basis of the subject-specific anatomy (52). Next, the activation per atlas region was averaged over grid points for each participant. Group-level activations were then calculated by averaging the activity over participants per atlas region (53). The early visual cortex ROI (EVC) corresponded to the ‘occipital pole’ parcels from the Destrieux atlas and the inferior temporal ROI (IT) was a combination of the ‘temporal lateral fusiform’, ‘temporal lateral’ and ‘temporal lateral and lingual’ parcels. Activation in the IT ROI was calculated by applying PCA to the three parcels and taking the first principal component. Because data are z-scored over time during PCA, to ensure that the activation in the EVC ROI was comparable to activation in IT, we also z-scored these data.

## Author contributions

Conceptualization: all authors.; Formal analysis: N.D., L.A.; Methodology: all authors; Project administration: N.D.; Resources: N.D., M.v.G.; Supervision: M.v.G.; Visualization: N.D.; Writing – original draft: N.D.; Writing – review & editing: all authors.

## Acknowledgements

The authors would like to thank Emma K. Ward for help with the linear mixed model statistics. This work is supported by VIDI grant (639.072.513) from the Netherlands Organization for Scientific Research.

## Competing interests

The authors declare no competing financial interests.

## Supplementary Note 1: Linear Mixed Model

**Table S1.** Model results. Bayesian information criterion (BIC) for linear mixed-effects model explaining imagery reactivation time with perception model time

| Model | Fixed effects | Random effects | df | BIC<sup>1</sup> |
| --- | --- | --- | --- | --- |
| 1 | Perception time | - | 3 | 323730.0 |
| 2 | Perception time | Subject | 4 | 323462.4 |
| 3 | Perception time | Subject + Perception time \| Subject | 6 | did not converge |
| Best fit | Perception time | Subject + Trial | 5 | 322911.5 |
| 5 | Perception time | Subject + Perception time \| Subject + Trial + Trial \| Perception time | 9 | did not converge |

## Supplementary Note 2: Permuted classifier

To exclude the possibility that the reversal in perception model activation during imagery was due to systematic biases in the data unrelated stimulus representations, we performed the same analysis with a classifier in which the labels of classes were permuted, removing information about the stimulus class. The reactivation time is plotted in Supplementary Figure S1 and the LMM parameters are plotted in Supplementary Table S2. The sequential reactivation disappeared when using shuffled classifiers as perception time did not significantly predict imagery reactivation time anymore (*t*(98160) = −0.762, *p* = 0.446).

**Fig. S1.**
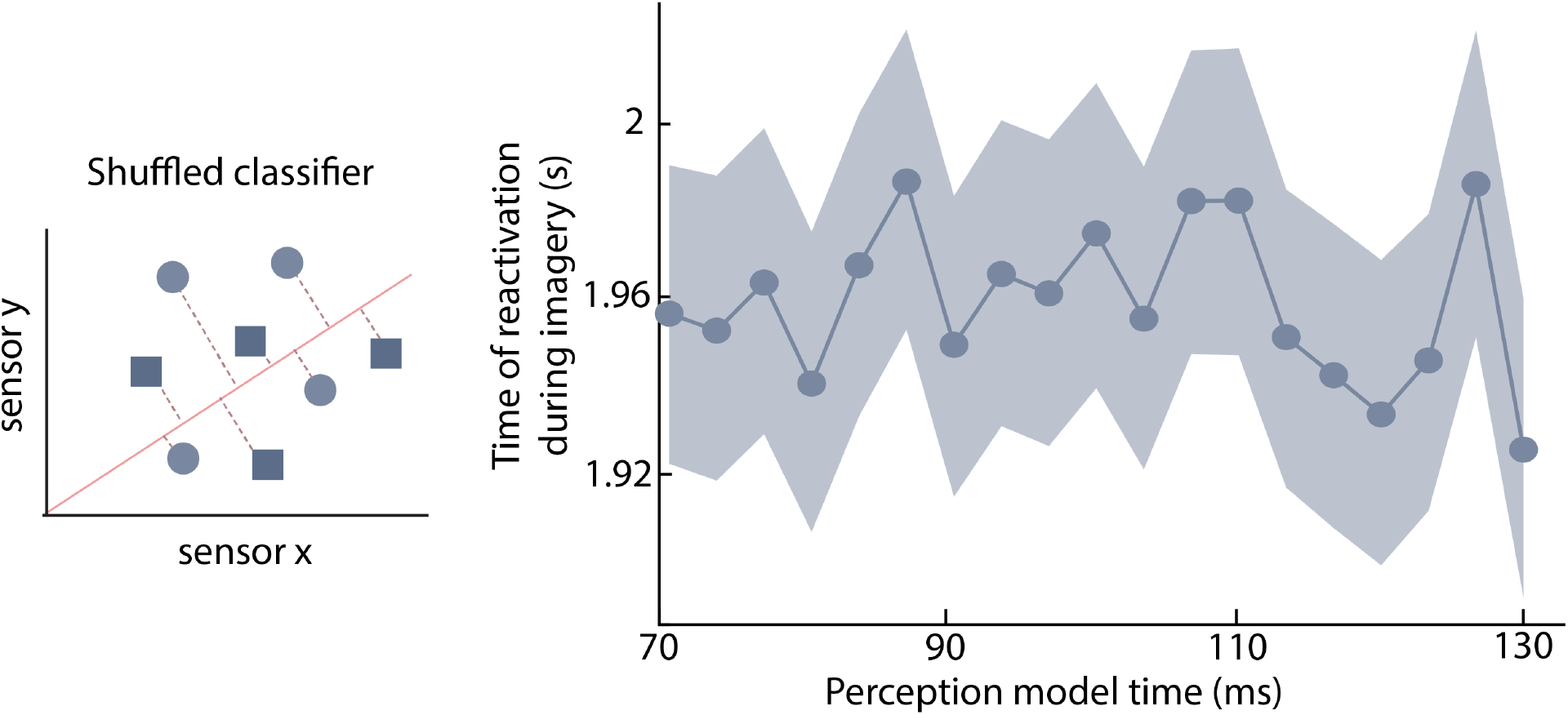
Reactivation timing results for shuffled classifier. To test whether the reactivation pattern was due to irrelevant structure in the data we used shuffled perception classifiers and performed the same reactivation analysis as before (Figure 3). No reactivation sequence is present using this classifier, confirming that the results are due to actual reactivation and not confounds in the structure of the data.

**Table S2.** Model results. Bayesian information criterion (BIC) for linear mixed-effects model explaining imagery reactivation time with perception model time

| Model | Fixed effects | Random effects | df | BIC |
| --- | --- | --- | --- | --- |
| 1 | Perception time | - | 3 | 326310.8 |
| 2 | Perception time | Subject | 4 | 326042.3 |
| 3 | Perception time | Subject + Perception time \| Subject | 6 | did not converge |
| Best fit | Perception time | Subject + Trial | 5 | 325415.1 |
| 5 | Perception time | Subject + Perception time \| Subject + Trial + Trial \| Perception time | 9 | did not converge |

## Supplementary Note 3: Effect low-pass filter

We removed high frequency noise from the distance values for the main analyses by applying a 30Hz low-pass filter. Qualitatively similar results were obtained without using the filter (see Supplementary Figure S2). There was still a significant main effect of perception time (*t*(98160) = −4.153, *p* = 0.0000328) where models of later perception time points were associated with earlier imagery reactivation times (*β* = −0.8942, SD = 0.2153) and the results of the shuffled classifier remained non-significant.

**Fig. S2.**
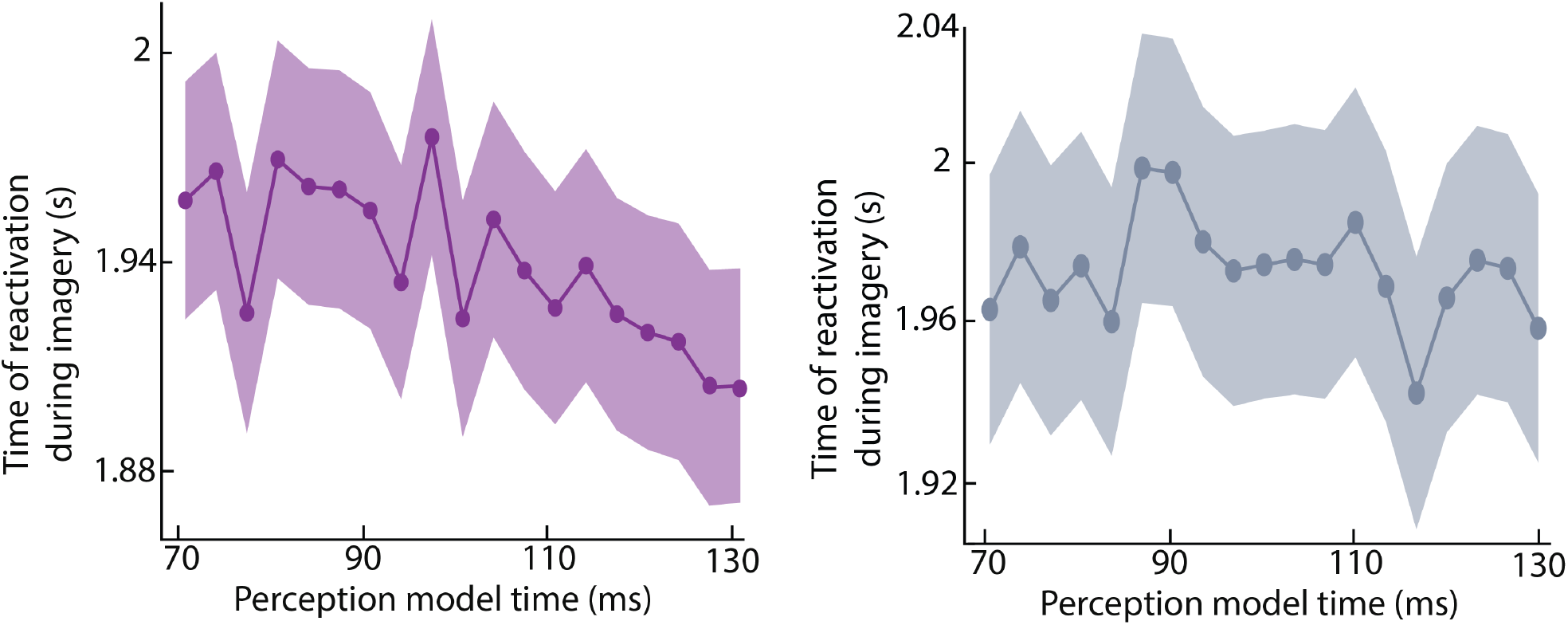
Reactivation timing results without low-pass filter. To test whether the reactivation pattern was somehow imposed by the low-pass filter, we performed the same analyses on the unfiltered data using the true classifier (left) and the permuted classifier (right). The results remained stable without the low-pass filter.

## Notes

http://hdl.handle.net/11633/di.dcc.DSC_2017.00072_245

## References

1. J. Patrick Card and Robert Y Moore. Organization of lateral geniculate-hypothalamic connections in the rat. Journal of Comparative Neurology, 284(1):135–147, 1989. ISSN 10969861. doi: 10.1002/cne.902840110.

2. R. Reid and Jose Manuel Alonso. Specificity of monosynaptic connections from thalamus to visual cortex. Nature, 378(6554):281–284, 1995. ISSN 00280836. doi: 10.1038/378281a0.

3. D. H. Hubel and T. N. Wiesel. Receptive fields and functional architecture of monkey striate cortex. Journal of Physiology, 195(1):215–243, mar 1968. ISSN 0022-3751. doi: papers://47831562-1F78-4B52-B52E-78BF7F97A700/Paper/p352.

4. J. Maunsell. Visual processing In monkey extrastriate cortex. Annual Review of Neuroscience, 10(1):363–401, 1987. ISSN 0147006X. doi: 10.1146/annurev.neuro.10.1.363.

5. S J Thorpe and M Fabre-Thorpe. Seeking categories in the brain. Science, 291(5502): 260–3, jan 2001. ISSN 0036-8075.

6. R Vogels and G A Orban. Coding of stimulus invariances by inferior temporal neurons. Progress in brain research, 112:195–211, 1996. ISSN 0079-6123.

7. K. Seeliger, M. Fritsche, U. Güçlü, S. Schoenmakers, J. M. Schoffelen, S. E. Bosch, and M. A.J. van Gerven. Convolutional neural network-based encoding and decoding of visual object recognition in space and time. NeuroImage, pages 1–14, 2017. ISSN 10959572. doi: 10.1016/j.neuroimage.2017.07.018.

8. S Thorpe, D Fize, and C Marlot. Speed of processing in the human visual system. Nature, 381(6582):520–522, 1996. ISSN 0028-0836. doi: 10.1038/381520a0.

9. The neural dynamics of face detection in the wild revealed by MVPA. The Journal of neuroscience, 34(3):846–54, jan 2014. ISSN 1529-2401. doi: 10.1523/jneurosci.3030-13.2014.

10. Less is more: Expectation sharpens representations in the primary visual cortex. Neuron, 75:265–270, 2012. ISSN 08966273. doi: 10.1016/j.neuron.2012.04.034.

11. Nadine Dijkstra, Sander E Bosch, and Marcel A.J. van Gerven. Shared neural mechanisms of visual perception and imagery. Trends in Cognitive Sciences, 23:18–29, 2019. ISSN 13646613. doi: 10.1016/j.tics.2019.02.004.

12. Vividness of visual imagery depends on the neural overlap with perception in visual areas. Journal of Neuroscience, 37(5):1367–1373, 2017. ISSN 0270-6474, 1529-2401. doi: 10.1523/JNEUROSCI.3022-16.2016.

13. Distributed neural systems for the generation of visual images. Neuron, 28:979–990, 2000. ISSN 08966273. doi: 10.1016/S0896-6273(00)00168-9.

14. Sue-Hyun Lee, Dwight J Kravitz, and Chris I Baker. Disentangling visual imagery and perception of real-world objects. NeuroImage, 59(4):4064–73, feb 2012. ISSN 1095-9572. doi: 10.1016/j.neuroimage.2011.10.055.

15. Leila Reddy, Naotsugu Tsuchiya, and Thomas Serre. Reading the mind’s eye: Decoding category information during mental imagery. NeuroImage, 50(2):818–825, apr 2010. ISSN 10538119. doi: 10.1016/j.neuroimage.2009.11.084.

16. Anke Marit Albers, Peter Kok, Ivan Toni, H. Chris Dijkerman, and Floris P De Lange. Shared representations for working memory and mental imagery in early visual cortex. Current Biology, 23:1427–1431, 2013. ISSN 09609822. doi: 10.1016/j.cub.2013.05.065.

17. Joel Pearson, Colin W G Clifford, and Frank Tong. The functional impact of mental imagery on conscious perception. Current Biology, 18(13):982–6, jul 2008. ISSN 0960-9822. doi: 10.1016/j.cub.2008.05.048.

18. Mario Senden, Thomas C. Emmerling, Rick van Hoof, Martin A. Frost, and Rainer Goebel. Reconstructing imagined letters from early visual cortex reveals tight topographic correspondence between visual mental imagery and perception. Brain Structure and Function, 224(3):1167–1183, 2019. ISSN 18632661. doi: 10.1007/s00429-019-01828-6.

19. Merav Ahissar and Shaul Hochstein. The reverse hierarchy theory of visual perceptual learning. Trends in Cognitive Sciences, 8(10):457–464, 2004. doi: 10.1016/j.tics.2004.08.011.

20. Shaul Hochsteinand Merav Ahissar. View from the Top: Hierarchies and reverse hierarchies in the visual system. Neuron, 36(5):791–804, 2002. doi: 10.1016/S0896-6273(02)01091-7.

21. Stephen M Kosslyn, Giorgio Ganis, and William L Thompson. Neural foundations of imagery. Nature Reviews Neuroscience, 2(9):635–642, 2001. ISSN 1471-003X. doi: 10.1038/35090055.

22. Joel Pearson and Rebecca Keogh. Redefining visual working memory: A cognitive-strategy, brain-region approach. Current Directions in Psychological Science, pages 266–273, apr 2019. ISSN 0963-7214. doi: 10.1177/0963721419835210.

23. Juan Linde-Domingo, Matthias S. Treder, Casper Kerren, and Maria Wimber. Evidence for a reversal of the neural information flow between object perception and object reconstruction from memory. Nature Communications, 10(1):300913, dec 2019. ISSN 0003-987X. doi: 10.1001/archderm.143.9.1131.

24. Eelke Spaak, Kei Watanabe, Shintaro Funahashi, and Mark G Stokes. Stable and dynamic coding for working memory in primate prefrontal cortex. The Journal of Neuroscience, 37 (27):6503–6516, 2017. doi: 10.1523/JNEUROSCI.3364-16.2017.

25. H Helmholtz. Physiological Optics, Vol. III: The Perceptions of Vision. Optical Society of America, Rochester, NY, 1925.

26. Karl Friston. A theory of cortical responses. Philosophical Transactions of the Royal Society of London B: Biological Sciences, 360(1456), 2005.

27. David C. Knill and Alexandre Pouget. The Bayesian brain: The role of uncertainty in neural coding and computation. Trends in Neurosciences, 27(12):712–719, 2004. doi: 10.1016/j.tins.2004.10.007.

28. Andre M. Bastos, W. Martin Usrey, Rick A. Adams, George R. Mangun, Pascal Fries, and Karl J. Friston. Canonical microcircuits for predictive coding. Neuron, 76(4):695–711, 2012. doi: 10.1016/j.neuron.2012.10.038.

29. Diego Lozano-Soldevilla and Rufin VanRullen. The Hidden Spatial Dimension of Alpha: 10-Hz Perceptual Echoes Propagate as Periodic Traveling Waves in the Human Brain. Cell Reports, 26(2):374–380.e4, 2019. doi: 10.1016/j.celrep.2018.12.058.

30. Nadine Dijkstra, Pim Mostert, Floris P. de Lange, Sander Bosch, and Marcel A. J. van Gerven. Differential temporal dynamics during visual imagery and perception. eLife, 7: 1–16, may 2018. ISSN 2050-084X. doi: 10.1101/226217.

31. L. Isik, E. M. Meyers, J. Z. Leibo, and T. Poggio. The dynamics of invariant object recognition in the human visual system. Journal of Neurophysiology, 111(1):91–102, jan 2014. ISSN 0022-3077. doi: 10.1152/jn.00394.2013.

32. An optimal oscillatory phase for pattern reactivation during memory retrieval. Current biology, 28(21):3383–3392.e6, nov 2018. ISSN 1879-0445. doi: 10.1016/j.cub.2018.08.065.

33. Finding decodable information that can be read out in behaviour. NeuroImage, 179:252–262, oct 2018. ISSN 1053-8119. doi: 10.1016/J.NEUROIMAGE.2018.06.022.

34. Victor A.F. Lamme and Pieter R. Roelfsema. The distinct modes of vision offered by feedforward and recurrent processing. Trends in Neurosciences, 23(11):571–579, 2000. doi: 10.1016/S0166-2236(00)01657-X.

35. Cyriel M.A. Pennartz, Shirin Dora, Lars Muckli, and Jeannette A.M. Lorteije. Towards a unified view on pathways and functions of neural recurrent processing. Trends in Neurosciences, 1528:1–15, aug 2019. ISSN 01662236. doi: 10.1016/j.tins.2019.07.005.

36. Daniel Kersten, Pascal Mamassian, and Alan Yuille. Object perception as Bayesian inference. Annual review of psychology, 55:271–304, 2004. doi: 10.1146/annurev.psych.55.090902.142005.

37. Samuel J Gershman. The generative adversarial brain. Technical report, 2019.

38. Rick Grush. The emulation theory of representation: Motor control, imagery, and perception. Behavioral and Brain Sciences, 27:377–442, 2004.

39. J.A. Hobson and K.J. Friston. Waking and dreaming consciousness: Neurobiological and functional considerations. Progress in Neurobiology, 98(1):82–98, jul 2012. ISSN 0301-0082. doi: 10.1016/J.PNEUROBIO.2012.05.003.

40. Samuel T Moulton and Stephen M Kosslyn. Imagining predictions: Mental imagery as mental emulation. Philosophical Transactions of the Royal Society B: Biological Sciences, 364(1521):1273–1280, 2009. ISSN 14712970. doi: 10.1098/rstb.2008.0314.

41. Stephen M Kosslyn and William L Thompson. When is early visual cortex activated during visual mental imagery? Psychological bulletin, 129(5):723–746, 2003. doi: 10.1037/0033-2909.129.5.723.

42. Stephenie A. Harrison and Frank Tong. Decoding reveals the contents of visual working memory in early visual areas. Nature, 458(7238):632–635, apr 2009. doi: 10.1038/nature07832.

43. Arjen Stolk, Ana Todorovic, Jan Mathijs Schoffelen, and Robert Oostenveld. Online and offline tools for head movement compensation in MEG. NeuroImage, 68:39–48, 2013. ISSN 10538119. doi: 10.1016/j.neuroimage.2012.11.047.

44. Robert Oostenveld, Pascal Fries, Eric Maris, and Jan Mathijs Schoffelen. FieldTrip: Open source software for advanced analysis of MEG, EEG, and invasive electrophysiological data. Computational Intelligence and Neuroscience, 2011, 2011. doi: 10.1155/2011/156869.

45. Pim Mostert, Peter Kok, and Floris P. de Lange. Dissociating sensory from decision processes in human perceptual decision making. Scientific Reports, 5:18253, dec 2015. ISSN 2045-2322. doi: 10.1038/srep18253.

46. N Huang, Z Shen, S Long, M Wu, H Shih, Q Zheng, N Yen, C Tung, and H Liu. The empirical mode decomposition and the Hilbert spectrum for nonlinear and non-stationary. Proceedings Mathematical, 454:903–995, 1998.

47. Gabriel Rilling, Patrick Flandrin, and Paulo Goncalves. On empirical mode decomposition and its algorithms. In IEEE-EURASIP workshop on nonlinear signal and image processing, volume 3, pages 8–11, 2003.

48. Gang Wang, Xian Yao Chen, Fang Li Qiao, Zhaohua Wu, and Norden E. Huang. On intrinsic mode function. Advances in Adaptive Data Analysis, 2(3):277–293, 2010. ISSN 17935369. doi: 10.1142/S1793536910000549.

49. Stefan Haufe, Frank Meinecke, Kai Görgen, Sven Dähne, John Dylan Haynes, Benjamin Blankertz, and Felix Bießmann. On the interpretation of weight vectors of linear models in multivariate neuroimaging. NeuroImage, 87:96–110, 2014. doi: 10.1016/j.neuroimage.2013.10.067.

50. B D Van Veen, W van Drongelen, M Yuchtman, and A Suzuki. Localization of brain electrical activity via linearly constrained minimum variance spatial filtering. IEEE transactions on biomedical engineering, 44:867–880, 1997. doi: 10.1109/10.623056.

51. Mariya E. Manahova, Pim Mostert, Peter Kok, Jan-Mathijs Schoffelen, and Floris P de Lange. Stimulus familiarity and expectation jointly modulate neural activity in the visual ventral stream. Journal of Cognitive Neuroscience, 30(9):1366–1377, sep 2018. ISSN 0898-929X. doi: 10.1162/jocn_a_01281.

52. Christophe Destrieux, Bruce Fischl, Anders Dale, and Eric Halgren. Automatic parcellation of human cortical gyri and sulci using standard anatomical nomenclature. NeuroImage, 53 (1):1–15, 2010. ISSN 10538119. doi: 10.1016/j.neuroimage.2010.06.010.

53. Marieke E van de Nieuwenhuijzen, Eva W P van den Borne, Ole Jensen, and Marcel A J van Gerven. Frontiers in Systems Neuroscience, page 42. ISSN 1662-5137. doi: 10.3389/fnsys.2016.00042.

